# Compound biological signatures facilitate phenotypic screening and target elucidation

**DOI:** 10.1101/041947

**Authors:** Alvaro Cortes Cabrera, Daniel Lucena-Agell, Mariano Redondo-Horcajo, Isabel Barasoain, Fernando Diaz, Bernhard Fasching, Paula Petrone

## Abstract

We developed a statistical method, BIOSEA, able to identify molecules that are capable of reproducing a desired cellular phenotype, by scanning a large compound collection based on biological similarity. Our method leverages highly incomplete and noisy compound bioactivity signatures from historical high-throughput screening campaigns. We applied our approach in a phenotypic screening workflow and found novel nanomolar inhibitors of cell division that reproduce the mode of action of reference natural products. In a drug discovery setting, our biological hit expansion protocol revealed new inhibitors of the NKCC1 co-transporter for autism spectrum disorders. Furthermore, we demonstrate BIOSEA’s capabilities to predict novel targets for old compounds. We report new activities for the drugs nimedipine, fluspirilene and pimozide applicable for compound repurposing and rationalizing drug side effects. Our results highlight the opportunities of reusing public bioactivity data for prospective drug discovery applications where the target or mode of action is not known.

---

The last two decades have witnessed a dramatic change in our understanding of the interactions between small molecules and biological systems^1^. We are currently experiencing an important shift from reductionism^2^, to a more pragmatic vision of living organisms as complex entities whose study cannot be effectively tackled by treating each part in isolation^3^. In this new framework, phenotypic screening has reemerged as a practical solution to finding novel chemical entities, which provide insights into the underlying cellular processes leading to new opportunities for drug discovery^4^. However, phenotypic screening brings about its own challenges such as the identification of the mechanism of action (MoA) and the effective target or targets responsible for these phenotypes^5^. Target and MoA identification play a fundamental role, paving the way to lead optimization via structural approaches, structure-activity relationships (SAR) rationalization and also by improving awareness about ADME (absorption, distribution, metabolism and excretion), safety liabilities and drug-drug interactions. The chemical biology and genomics fields have stood up to the challenge producing elegant solutions for these problems^6^, such as CRISPR/Cas9 gene editing^7^ and chemical proteomics-based techniques^8^ that yet, bear inherent limitations such as chemical modification of the compounds, challenging specificity or high costs^8^. In the light of these advances, novel computational efforts are now in order to support phenotypic screening. In this work, we demonstrate how to leverage historical bioassay data as compound biological signatures to facilitate phenotypic deconvolution tasks, such as target identification and phenotypic library design. We report here a novel target and chemical structure-independent screening technique to find new chemical compounds to modulate a desired phenotype in a drug discovery setting.

Large amounts of pharmacological and bioactivity data accumulate both in public repositories and in proprietary databases in the pharmaceutical industry. A few approaches have attempted to use historical compound annotation data for target identification. For example, the TarFishDock^9^, TargetHunter^10^, Similarity Ensemble Approach^11,12^, ChemMapper^13^ and HitPick^14^ methods, rely on either 2D or 3D chemical information of ligands or targets and an established molecular structure similarity metric. These methods have been exploited with success in tasks such as the prediction of targets responsible for drug side effects^15^. However, an important limitation is the fact that these methods depend on the availability of crystal structures for the target, or the structural similarity between the compounds and previously known target ligands which confines their performance to well-explored areas in chemical space.

In recent years, it has been demonstrated that the cellular phenotype of compounds can be described, and even predicted, by a compound’s biological signature: an array of numbers representing the perturbation caused by a chemical entity of interest measured throughout several cellular and biochemical assays^14^. In this context, compounds sharing a similar biological profile tend to modulate similar targets, without necessarily having similar chemical structure^16^. In the past, we and others^17-22^ have proposed the use of historical high-throughput screening (HTS) activity data to represent molecules in the biological space. The so-called HTS fingerprints (HTSFP) have proven successful for the identification of new targets for known compounds^22^. Since they are structure-independent, HTSFP enable scaffold-independent linking of drug-like molecules to less frequently explored regions in chemical space such as those of complex natural products^16,22^.

By integrating external pharmacological sources of information and HTS data, we have implemented BIOSEA - Biological Similarity Ensemble Approach - a novel statistical classifier that is able to establish new relationships between targets and small molecules on the basis of the mutual information between compound biological signatures. Algorithmically, BIOSEA was inspired by the method developed by Pearson^23^ or sequence similarity searches (BLAST) and the Similarity Ensemble Approach (SEA) developed by Keiser et al.^11^ to compare chemical structures.

In this work, first we report an unprecedented phenotypic screening workflow in which BIOSEA was applied to find compounds that modulate a desired cellular phenotype across large molecular libraries. We validated this procedure in two projects: (1) the discovery of drug-like inhibitors of cell division and (2) the identification of novel chemical entities to alter intracellular ion concentration by inhibiting the Na^+^ K^+^2Cl^−^ co-transporter (NKCC1) for the indication of autism. In a following section, we demonstrate the application of BIOSEA for target identification, validated with both retrospective data and prospective experiments that shed light on the polypharmacology of known drugs. Consistently throughout these applications, BIOSEA revealed new relationships between molecules and targets, which are difficult to detect even by a trained chemist’s eye.

## RESULTS

### BIOSEA identifies molecules that reproduce a desired cellular phenotype

BIOSEA enables phenotypic screening without the need of information about the chemical structure of the target or associated ligands (Fig. 1a). This capability was used in two drug discovery workflows to find novel chemotypes that: (i) inhibit cell division and (ii) block NKCC1 mediated modulation of intracellular ion concentration. In the first scenario, the cellular phenotype was characterized, and both target and MoA of the proposed compounds were identified using cellular and biochemical assays. In the second scenario, we applied BIOSEA for biological hit expansion with a NKCC1 cellular assay and compared its performance to standard methods that rely on chemical structure.

**Figure 1.**
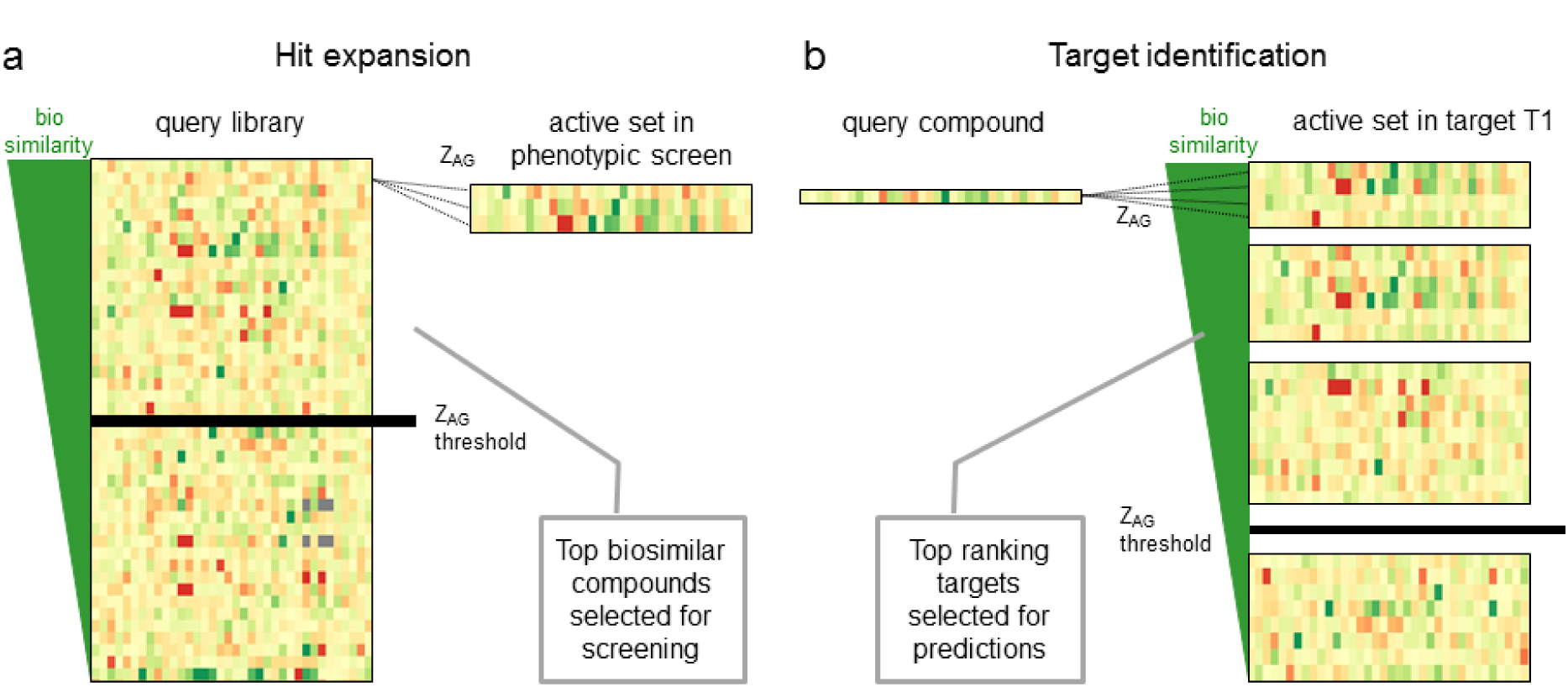
(a) For virtual screening, BIOSEA calculates the aggregated biological similarity (Z_AG_) of each compound in a library against a set of biological signatures from HTS (HTSFP) corresponding to compounds that are active in a cellular assay. The query library is sorted by Z_AG_ and the top ranking compounds are selected for a confirmatory screen. (b) For target identification, BIOSEA calculates the Z_AG_ of a query compound against a list of HTSFP sets T_1_…T_N_. Each fingerprint set represents the compounds that are active in a given target. Target sets are ranked according to Z_AG_ and the top 10 targets are defined as target predictions for the query compound.

### Phenotypic screening with BIOSEA: inhibition of cell division

As a reference set, we have selected six well-known natural products that perturb cell division in two different ways: paclitaxel acts as a microtubule-stabilizing agent while the remaining compounds have an inhibitory effect on tubulin polymerization. Virtual screening using these reference compounds and a library of circa 365,000 compounds was performed to predict 20 biologically similar compounds that would reproduce the phenotype of cell cycle arrest.

In a first validation step, using a cell cycle arrest screen in A549 cells, we could observe that 5 out 20 compounds were able to produce visible effects (rounding of cells) at a concentration of 5 μM which was indicative of a cytostatic behavior. The cytotoxic activity of the positive compounds (hereby named CT1 to CT5) was verified in the same cell line, with most IC_50_ values in the nanomolar range (Table 1). Additionally, it was confirmed that all 5 compounds completely depolymerized the interphasic cellular microtubule network (Fig. 2a). To establish tubulin as the target that modulated microtubule dynamics, the time-course of the assembly of tubulin was recorded in excess concentration of the hit compounds (Fig. 2b). All compounds were found to inhibit tubulin polymerization *in vitro* and in cells. Binding of active compounds in known tubulin depolymerizing binding sites (i.e. colchicine, vinca, maytansine, and eribulin) was investigated using various competition assays (Fig. 2c, Supplementary Fig. 1 and Supplementary Fig. 2). CT5 was found to bind to the eribulin biding site. The low solubility and weak activity of CT2 prevented any competition assay. CT1, CT3 and CT4, were confirmed to bind to the colchicine site and a pharmacophore model was proposed on the basis of colchicine-tubulin crystallographic information^24^ to rationalize the binding mode of these structurally diverse chemotypes (Fig. 2d, Supplementary Information). Although the compounds do not share the same scaffold, the model showed a clear superimposition of the di-and tri-methoxyphenyl moieties and good shape and chemical complementarity with the reference compound, which indicates a similar binding pattern.

**Figure 2.**
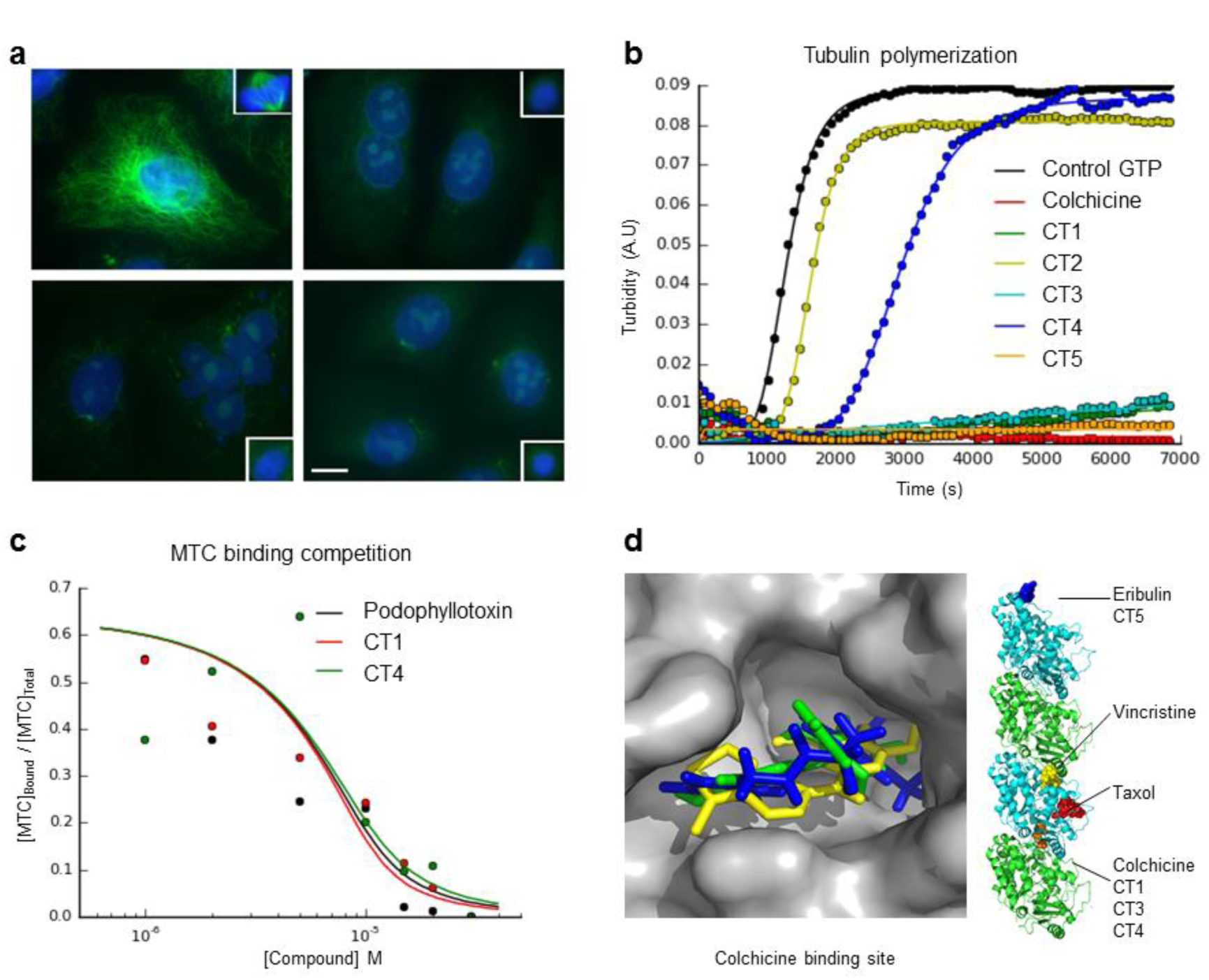
(a) Upon treatment with the compounds, the A549 cells showed type IV aberrant mitotic spindles^52^ consisting of a ball of condensed chromosomes and no microtubules. Upper-left: Control; upper-right: colchicine effects at 5 μM; lower-left: CT3 effects at 5 μM; and lower-right: CT5 effects at 25 μM (b)Time-course tubulin polymerization. (c) Displacement curves for the MTC probe in the colchicine binding pocket for compounds CT1, CT4 and podophyllotoxin. (d) Structural model of colchicine-site binders and summary of tubulin binding pockets for the active compounds (Colchicine in green, CT3 in yellow and CT4 in blue).

**Table 1:**
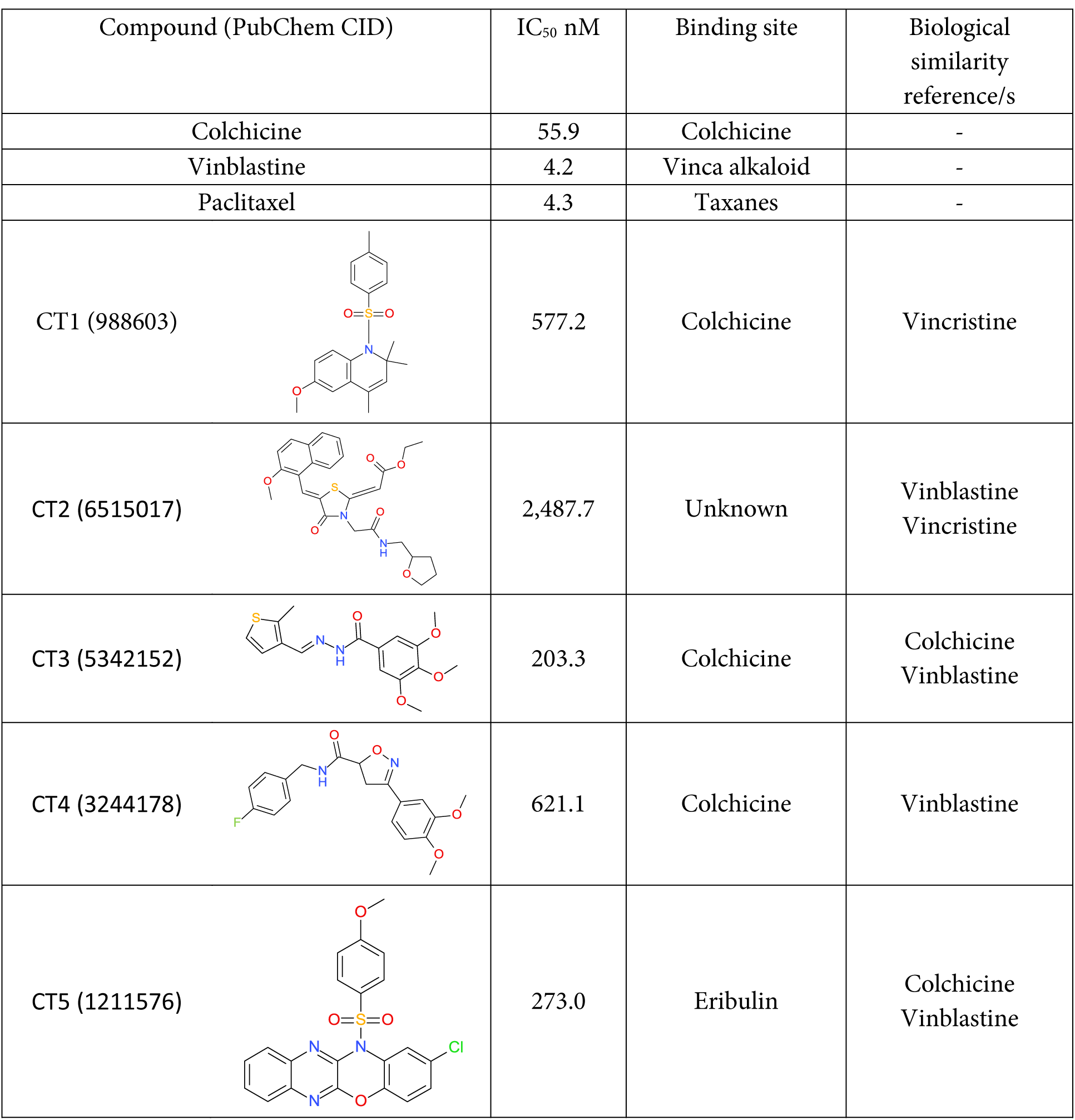
Phenotypic screening reference compounds (colchicine, vinblastine and paclitaxel) and cell arrest hits (CT1-CT5).

| Compound (PubChem CID) | IC<sub>50</sub> nM | Binding site | Biological similarity reference/s |
| --- | --- | --- | --- |
| Colchicine | 55.9 | Colchicine | - |
| Vinblastine | 4.2 | Vinca alkaloid | - |
| Paclitaxel | 4.3 | Taxanes | - |
| CT1 (988603) | 577.2 | Colchicine | Vincristine |
| CT2 (6515017) | 2,487.7 | Unknown | Vinblastine<br>Vincristine |
| CT3 (5342152) | 203.3 | Colchicine | Colchicine<br>Vinblastine |
| CT4 (3244178) | 621.1 | Colchicine | Vinblastine |
| CT5 (1211576) | 273.0 | Eribulin | Colchicine<br>Vinblastine |

To deconvolute the information encoded in the HTSFP that leads to successful target-specific compound predictions, we evaluated the individual contribution of every HTSFP assay to the global biological similarity to the reference compounds. Thus, for each of the 5 hits, the top 10 HTSFP components contributing most to the similarity were identified by a leave-one-out procedure (Supplementary Fig. 3, Methods) and classified into biochemical or cell-based assays (Supporting Table 1). Even though mutual information is a metric that is calculated based on the full range of the HTSFP fingerprint, this approach provided the following observations on these highly informative assays (HIA): **a) there is no unique pattern for HIA that is common for all active compounds**; instead, each compound attributes its biological similarity to a different part of the fingerprint. For example, two compounds that were found to bind in the colchicine pocket, CT1 and CT4, do not have any HIA in common (Supplementary Fig. 3), and their biologically closest reference compounds are also different (vincristine and vinblastine, respectively); **b) no HIA exists for the phenotype of interest (i.e. cell cycle arrest);** only 2 assays measure unspecific cytotoxicity endpoints for *Giardia lamblia* and *Escherichia coli* (Supplementary Table 1). The remaining HIA cover a diverse range of biochemical and cell-based assays for targets such as the sentrin-specific protease 7, the sialic acid acetylesterase or the vainilloid receptor 1, which are not associated to microtubule formation. It is important to note that there is no tubulin-related assay in HTSFP; **c) cell-based and biochemical assays seem to contribute alike although most compounds resulted inactive in these HIA**. While compound activity in cell-based assays (Z-Score>2) may provide information on general cellular perturbations (e.g. cytotoxicity, signaling pathways, etc.), inactivity throughout biochemical or cellular assays (|Z-Score|<2) may bring information relevant for ruling out competing mechanisms leading to a given phenotype. As an example, among its HIA the molecule CT2 has resulted inactive for procaspases 3 and 7-two biochemical assays-and inactive in a bacterial cell death screen in *E-coli*.

### Hit expansion on NKCC1

The primary goal of hit expansion efforts following an HTS campaign is to leverage informative lead molecules to rescue other active compounds that were neglected in the hit triage or missing in the primary screening library. In a phenotypic screen, this approach is specially challenging due to the diversity in chemical leads which may modulate several unknown targets - a context in which the chemical similarity principle for hit expansion may not apply.

The Roche NKCC1 inhibitor program aimed at correcting imbalances in the GABA developmental switch^25^, which controls the intracellular chloride concentration and shifts the functional role of GABA between neonatal and adult stages from excitatory to inhibitory. Mediated by the change in expression of two major ion co-transporters NKCC1 and KCC2, alterations in this mechanism possibly result in CNS disorders including epilepsy, autism spectrum disorder, schizophrenia and effects of traumatic brain injury^26^. The goal of the project was to identify NKCC1 inhibitors with high selectivity over peripherally expressed NKCCs and largely improved brain exposure compared to available compounds from the loop diuretic class such as bumetanide.

To this end, a primary screen of more than 1.2 million compounds within the Roche library was conducted. The screen had a low hit rate of 0.06% confirmed active compounds covering several compound classes. A standard approach that relies on the generation of 3D shapes and surface charge models (ROCS^27^) was applied and 430 compounds pharmacophorically similar to selected active compounds were submitted for profiling. This procedure resulted in a single novel hit with an affinity of 9 μM. For biological hit expansion using BIOSEA, 12 confirmed hits with IC_50_ in the low micromolar to submicromolar range for NKCC1 were used as the reference set for the virtual screening of the Roche 1.2-million-compound library. Compounds were sorted by biological similarity (i.e. e-value) to the reference set and the top 44 compounds were tested for their NKCC1 activity in the profiling cellular assay. This resulted in the identification of three molecules with an IC_50_ below 30 μM (Fig. 3). Not only was there a significant improvement in the hit rate by using BIOSEA compared to traditional approaches based on 2D or 3D chemical information^28^, but, in addition, two of these molecules provided novel chemotypes (Fig. 3) useful for further optimization efforts. The remaining hit had a very similar chemotype to the reference compounds, which served as a control result given that BIOSEA operates independently from chemical structure.

**Figure 3.**
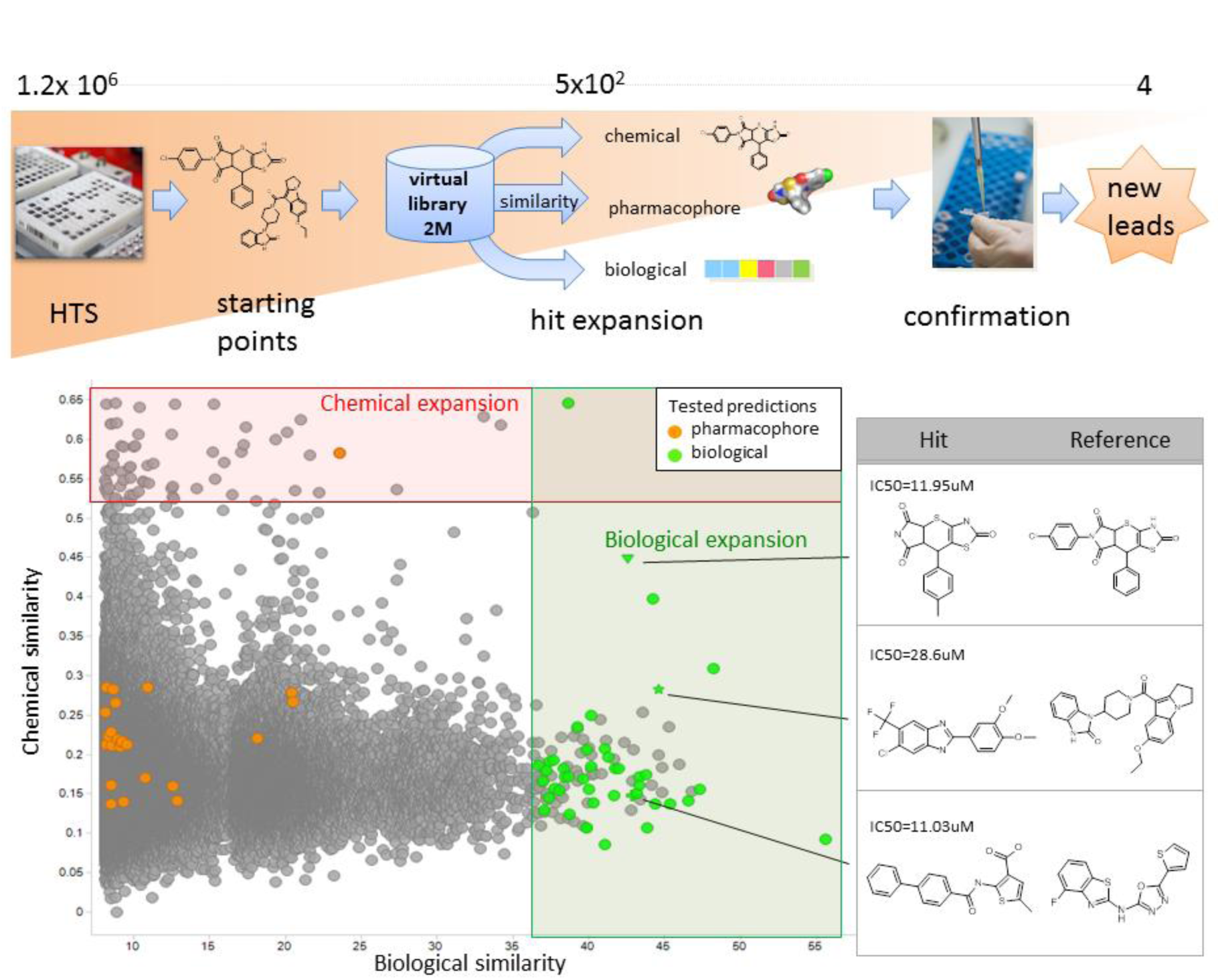
(Top) Phenotypic screening workflow for NKCC1. The primary screen of a 1.2 million compounds provided starting points for hit expansion within a 2-million-compound virtual library. Hit expansion was carried out by three complementary approaches: chemical, pharmacophore and biological similarity which produced in total ~500 compounds to be tested in a confirmatory dose-response assay, resulting in 4 leads with IC50<30 μM. (Bottom) Chemical similarity (Tanimoto distance of ECFP4 fingerprints) vs. biological similarity (represented by). The regions relevant for chemical and biological expansion are highlighted in red and green respectively. Green and orange dots represent NKCC1 tested biological and pharmacophore similarity predictions.

### BIOSEA can identify new targets for old compounds

Using BIOSEA, we investigated the polypharmacology of known drugs and evaluated the method’s potential to identify novel targets following the procedure described in Fig. 1b.

Our efforts were focused on 711 FDA-approved, withdrawn or experimental drugs. BIOSEA was used to assess their possible associations with public domain targets from the ChEMBL database^29^. Each molecule was assigned a maximum of 10 predicted targets above a minimum threshold of e-value, building a pool of 1,124 predictions, out of which 407 could be either experimentally validated or found in the literature. None of the reported target-compound associations were part of the training set and none of the attributed targets were part of the HTS fingerprint. Noteworthy, we predicted and experimentally confirmed 5 novel and unreported compound-target associations for drugs: pimozide, fluspirilene, reserpine, and nimodipine (Table 2). These examples show how this technology can provide: (i) deeper insights into the compound’s pharmacological profile responsible for its MoA, (ii) a rationale for previously identified side effects and (iii) opportunities for drug repurposing. **(i) Polypharmacology of pimozide and fluspirilene:** Pimozide and fluspirilene (Table 2) are antipsychotic drugs belonging to the diphenylbutylpiperidines (DPBPs) class, used conventionally for the treatment of schizophrenia and anxiety disorders. The established MoA of DPBPs is attributed to their potent blockade of the dopamine D_2_ receptors. Using BIOSEA, we correctly predicted activities against 7 other targets reported in the literature of which the serotonin transporter (SERT) and dopamine D_3_ receptor are common to both compounds (Supplementary Data Set 1). Previously unreported norepinephrine transporter (NET) activity for both pimozide and fluspirilene (IC_50_ = 110 nM, 480 nM respectively) was predicted and *in vitro* confirmed. NET is a well-known target for multiple mood and behavioral disorders such as depression, anxiety or attention deficit hyperactivity. Given a pimozide plasma concentration of 500 nM (recorded *in vivo* studies^30^), our reported submicromolar DPBPs activity in the NET is clinically relevant and contributes to the unique pharmacological profile of DPBPs in their treatment of intricate imbalances of the central nervous system. In particular, NET modulation may confer DPBPs with stimulant properties that are advantageous in the treatment of the negative symptoms of schizophrenia (e.g. emotional withdrawal). Additionally, BIOSEA was able to broadly capture the pharmacological profile of other antipsychotics (i.e. prochlorperazine and perphenazine), for which it could detect four or more annotations (Supplementary Data Set 1). The efficacy of those drugs in the treatment of schizophrenia and other mental illnesses has been previously linked to their ability to modulate multiple targets^31^ predominantly receptors subtypes and transporters of neurotransmitters (adrenaline, dopamine and serotonin). **(ii) Target identification provides a rationale for side effects:** In addition to NET, fluspirilene activity on the adrenergic α_2B_ receptor (ADRA2B, IC_50_ = 95 nM), a target related to vasoconstriction and blood pressure regulation^32^ was predicted and experimentally confirmed. Reducing blood pressure in patients with hypertension is documented^33^ as a side effect of fluspirilene, to which this ADRA2B activity could contribute. **(iii) Compound repurposing opportunity:** we predicted and experimentally confirmed the glucocorticoid receptor (GR) activity (IC_50_ = 640 nM) for the neuroprotective agent nimodipine. Established as a selective L-type voltage-dependent calcium channel antagonist, nimodipine is often used in the treatment of subarachnoid hemorrhage^34^. Animal studies have shown that nimodipine is able to improve cognitive performance^35^, block memory destabilization and prevent memory reconsolidation^36^. On the other hand, it is well established that glucocorticoids influence cognitive performance and have impairing effects on memory function and consolidation^37^. Post-traumatic stress disorder (PTSD) is reported to be associated with increased glucocorticoid receptor sensitivity^38^. Our finding, therefore, suggests opportunities in using nimodipine in connection with psychological disorders associated with GR such as PTSD, although this possibility should be studied in more detail.

**Table 2:**
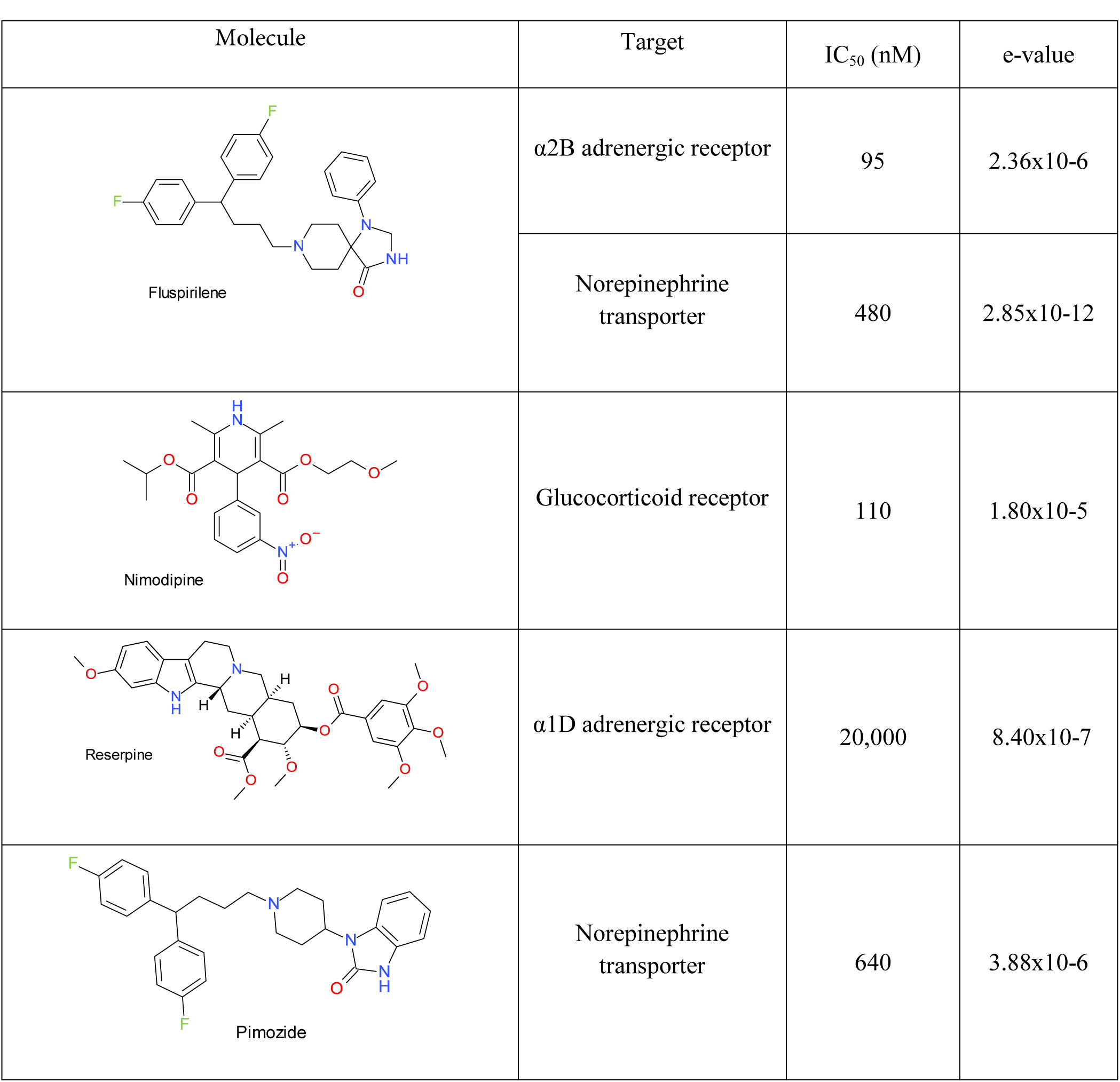
Novel targets for old drugs. Experimental conformation of the target identification predictions for pimozide, fluspirilene, reserpine, and nimodipine. Chemical structures, novel targets and experimental activities are included.

| Molecule | Target | IC<sub>50</sub> (nM) | e-value |
| --- | --- | --- | --- |
| Fluspirilene | α2B adrenergic receptor | 95 | 2.36x10<sup>-6</sup> |
|  | Norepinephrine transporter | 480 | 2.85x10<sup>-12</sup> |
| Nimodipine | Glucocorticoid receptor | 110 | 1.80x10<sup>-5</sup> |
| Reserpine | α1D adrenergic receptor | 20,000 | 8.40x10<sup>-7</sup> |
| Pimozide | Norepinephrine transporter | 640 | 3.88x10<sup>-6</sup> |

Taken together, 31% of validated target-compound relationships could be either positively confirmed, or putatively confirmed when a positive result was found in the literature for an orthologue target or convincing evidence of the relationship has been reported but is not quantified (Fig. 4a). Among compounds with validated predictions, 70 out of 193 were successfully assigned to at least one confirmed or putative target using BIOSEA. The same target identification benchmarking protocol was repeated using an in-house version of SEA^12^, which is based on chemical fingerprints (Supplementary Information). Only 4% of BIOSEA’s total predictions were shared with SEA, which hints at a high degree of complementarity between both methodologies. However, while approximately 80% of the positive results from SEA can be explained by the high structural similarity between the query compounds and the training set, only 32% of BIOSEA’s confirmed predictions could have been inferred in this way (Fig. 4c). The remaining predictions are not structurally obvious and suggest that HTSFP encode information that goes beyond a pure pharmacophore description and captures more general biological phenomena (see examples Fig. 4d).

**Figure 4.**
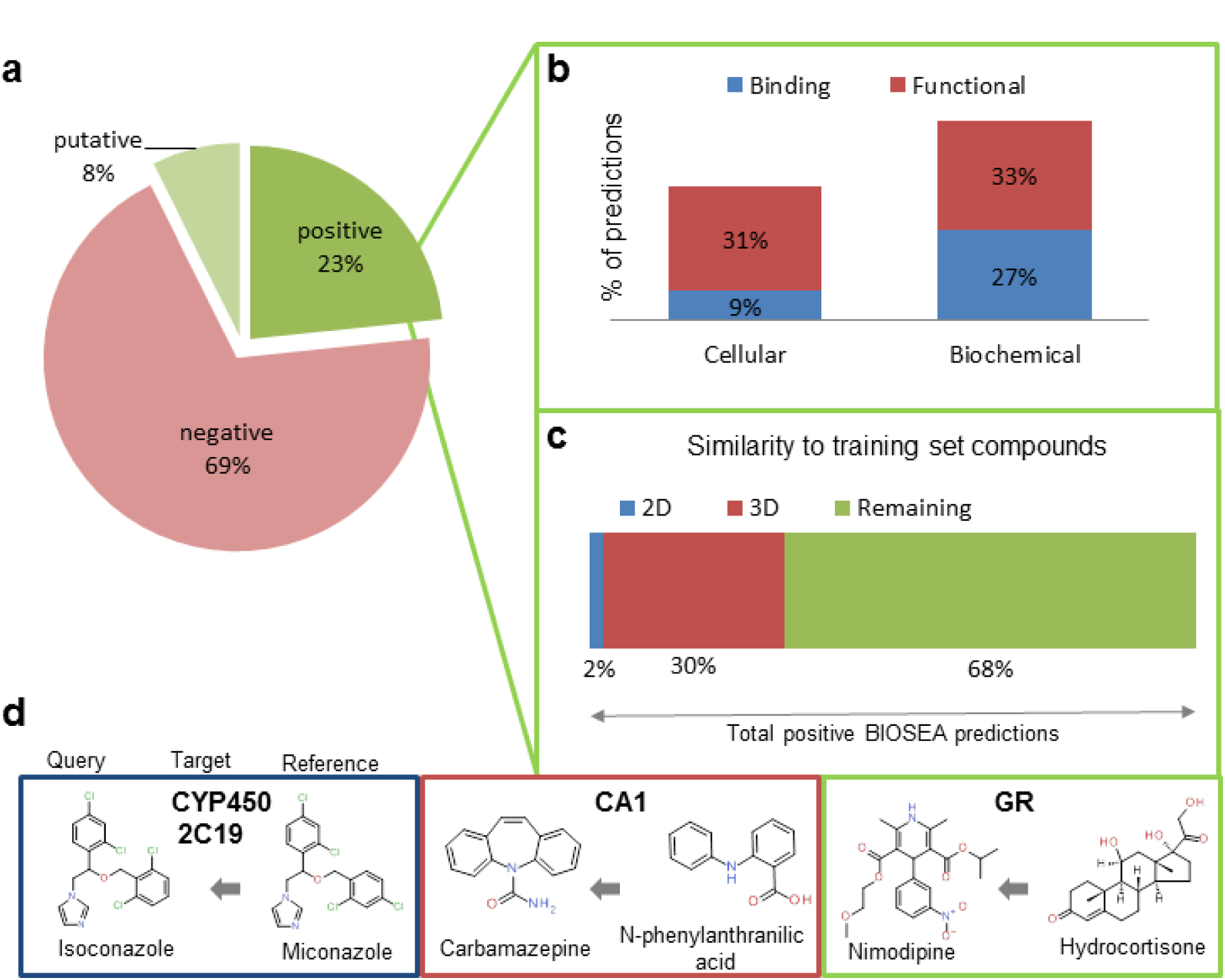
(a) Performance of BIOSEA in terms of verified predictions. (b) Categorization of positive results by types of assay. Assays were labeled as either functional (e.g. assay description provides a functional end-point for the compound’s effect), or binding (e.g. evidence for ligand binding); and biochemical or cell-based if the assay included only the target in an *in vitro* setup, or the whole cell in a more physiologic environment, respectively. (c) Biological similarity goes beyond 2D and 3D similarity. (d) Examples of similarity between predicted and reference molecules. From left to right: 2D similarity, 3D similarity and chemically unrelated (e) Previously unreported results of the target identification protocol for pimozide, fluspirilene, reserpine, and nimodipine.

Positive predictions from BIOSEA span both functional assays (63%) and binding assays (37%) (Fig. 4b). These percentages are in agreement with the biases in our database regarding the ratio of binding vs. functional annotations, indicating that the method performs equally well independently to the type of readout by which these predictions are confirmed. This result is consistent with previous observations making use of HTSFP which contain both cell-based and biochemical HTS assays^16,22^.

An interesting example among the literature-confirmed predictions is the inhibitory activity of amoxicillin (IC_50_ = 3.47 μM) on carbonic anhydrase XII^39^. This prediction illustrates the ability of BIOSEA to capture bioisosterism, since it was mostly based in the biologically similarity to compounds in the training set such as 4-sulfamoylbenzoic acid, sulpiride or indapamide, all of them sulfonamides. Though structurally unrelated to amoxicillin, all these compounds have in common a chemical group that is able to interact with the zinc atom in the enzyme’s active site (carboxylate or sulfonamide moieties respectively).

## DISCUSSION

Our results demonstrate that BIOSEA is able to classify compounds according to their biological signature from HTS data. Applied to phenotypic screening, this approach can identify compounds that are likely to produce a desired phenotype as captured in the biological signature of several reference compounds. We show that this procedure is possible even in the absence of information on the particular target(s) responsible for the compounds’ mode of action. The results for tubulin and NKCC1 show that the applicability domain of the method goes beyond the target, target families and phenotypes that have been historically screened and represented in the HTSFP. HTSFP are unique in the fact that they capture information on compound inactivity as well as compound activity, both equally relevant to the mutual information between fingerprints. While the desired phenotype, cell cycle arrest in this case, can be triggered by multiple mechanisms of action, interestingly, our hits bind to the same target and follow the mechanism of action of their reference compounds. This suggests that HTSFP are able to encode target-specific information (i.e. tubulin binding). The opportunity of using this approach for finding compounds that can reverse a particular disease phenotype remains an exciting avenue for further exploration.

In addition, the NKCC1 results suggest that the success of the screening effort can be evaluated not only in terms of hit enrichment but also in the novelty and variety of active chemical structures. For this reason, BIOSEA becomes an effective tool for building screening libraries for drug discovery programs, even if the target is novel and challenging, as is the case for NKCC1.

In target identification, our method can be used to discriminate among different drug targets by the similarity in the biological signatures of their corresponding ligands. Using our protocol we have explored the pharmacological profile of known drugs. Despite the fact that most of them have been optimized for selectivity, BIOSEA was able to accurately predict drug polypharmacology in many G-protein coupled receptors and a wide variety of enzyme classes, consistent with the annotation biases in public^40^ and industrial databases^41^ towards specific target families. In particular, we demonstrate the applicability of our method for drug repurposing and rationalization of drug side effects. Moreover, the majority of the confirmed predictions cannot be rooted in obvious chemical similarity to reference compounds. This shows that BIOSEA provides complementary information to traditional computational methodologies.

Despite its advantages, the approach has some inherent limitations, such as the availability of the HTS signature for a particular compound and the existence of suitable reference compounds for a given target or phenotype of interest. Additionally, for target identification, BIOSEA restricts its findings to the target domain that is known and has several annotated ligands. Nonetheless, with its performance validated in these in-house drug discovery and target identification scenarios, BIOSEA, offers a complementary and alternative source for hypothesis generation by capitalizing on already existing information resources.

Trained solely on public data, our innovative approach is within the reach of both academic and industrial research groups. We invite the scientific community at large to contribute and explore this technology by providing the computer code and the HTSFP public data set for download (https://github.com/accsc/BIOSEA), in the hope that it will foster present and future drug discovery efforts.

## METHODS

### Biological fingerprints from HTS

HTS fingerprints (HTSFP) were built using screening data from PubChem, consisting of 95 primary screening assays over a library of 365,231 molecules in a similar way to what was described in earlier works^21^ (Supplementary Information). HTS results were normalized using the average response and the standard deviation to transform all the readouts to Z-Score values. HTSFP-Roche were built in a similar fashion using Roche proprietary assay data.

#### Biosimilarity metric

A derived version of the mutual information (*I*) was introduced to estimate the similarity between two biological fingerprints Z_NMI_:

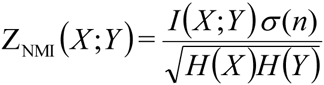

where X and Y are the HTSFP reduced to the assays in common for compounds x and y; *I*(*X*;*Y*) is the estimated mutual information using the Kraskov entropy estimator from k-nearest neighbor distances^42^; *H*(*X*) and *H*(*Y*) are the Shannon entropies^43^; and σ(n) was introduced to correct for the dependency in fingerprint-size *n*, parametrized based on a background distribution of random fingerprint comparisons (a more detailed procedure can be found in Supplementary Information). An optimal value of *k* = 10 was selected for the calculation of *I*(*X*;*Y*) based on previous parameter analysis (data not shown) and as trade-off between minimal fingerprint length required and a reliable Z_NMI_ calculation.

#### BIOSEA

A statistical method to classify compounds into target classes using biological signatures from HTS was developed, inspired by the BLAST^23^ and SEA^11^ algorithms. BIOSEA provides an e-value that represents the likelihood of a compound to be related by chance to a given compounds set, by aggregating the biological similarities (Z_NMI_) between the compound and the members of the set, and comparing the aggregated value to a random distribution of sums. To build the background distribution, random lists of HTSFP were generated with sizes in the range *s = 1* to *s = 100*. The aggregated biological similarity between every two pairs of lists of compounds with sizes *1* and *m* respectively was calculated. The average and standard deviation of the random sums of Z_NMI_ per number of comparisons (*l x m*) were calculated and fitted to two power-law equations (Supplementary Table 2). Aggregated biological similarities were transformed to Z-Scores (Z_AG_) and plotted in a histogram that conformed to an Extreme Value Distribution (EVD), whose location, scale and shape parameters were estimated (Supplementary Table 3). This distribution was used as a background for probability calculation and its conversion to an expectation value (e-value) that evaluates the significance of the relationships between given sets of fingerprints. The choice of Z_NMI_ threshold between fingerprints selects which compounds are relevant for the aggregated biological similarity and has an effect on the Z_AG_ distribution. Z_NMI_ values of 0.5 and 4.5 in increments of 0.1 were tested and minimal (Z_NMI_ = 2.6) and optimal values (Z_NMI_ = 4.0) were obtained. More detailed information can be found on Supplementary Information.

#### Individual assay contributions to the aggregated similarity

For each compound with tubulin depolymerizing activity, we carried out a leave-one-out procedure in which the aggregated biological similarity of an individual compound to the reference compounds (Z_AG_) was calculated after removing one assay at a time. The difference in Z_AG_ between using the complete fingerprint and the version with a removed component was computed and used to rank the assays. The first 10 assays with the highest similarity difference were selected for analysis.

#### Target identification database

For the training of BIOSEA, a database of 265 targets and their known ligands was built using dose-response annotations (IC_50_, EC_50_, AC_50_, K_i_, K_d_, Potency) from ChEMBLdb release 20^29^. Molecules with at least one measurement below 5 μM and none higher than 15 μM were considered active. Only targets with at least 10 annotated ligands were included. Frequent binders (actives in more than 5 assays) were removed from the datasets. A total of 21,157 annotations for 19,808 molecules were collected as the training set for target identification (Supplementary Data Set 2).

#### Predictions for target identification

711 known drugs were extracted from DrugBank^44^ and ChEMBLdb release 20. Predictions were carried out by BIOSEA trained on the target identification database. The compound biosimilarity threshold was defined as Z_NMI_>2.6. The threshold for compound-target predictions was e-value > 2.72x10^−5^ (equivalent to Z_AG_ > 8.0) resulting in a total pool of 1,124 predictions. The 164 predictions for which the target is represented in the HTSFP were eliminated from the statistics. A total of 407 remaining predictions were literature or experimentally verified. ChEMBLdb release 20, DrugBank and Thomsons Reuters Integrity databases were scanned for activity confirmation. PubMed and EMBASE databases were inspected with the text mining Linguamatics I2E tool for additional evidence. Predictions were confirmed as active by a retrieved dose-response value below a threshold of 20 μM, or they were supported by literature annotations without a numeric readout. A subset of 169 unreported testable hypotheses (accessible compounds and available assays, see Supplementary Data Set 1) was extracted for experimental evaluation. Analysis of the 2D compound similarity was performed using ECFP4 fingerprints generated by Pipeline Pilot 9.2 with a Tanimoto coefficient threshold of 0.8. 3D compound similarity was calculated using a ROCS-clone^27,45^ based on Gaussian shape similarity and a Tanimoto combo threshold of 1.5.

#### NKCC1 assay

HEK293 cells conditionally expressing NKCC1 were grown under doxycycline selection. Cell cultures were maintained at 37 °C in the presence of 5% CO_2_ and 85% of humidity. A volume of 50 μl containing 25,000 cells was plated on poly-D-lysine dishes and induced for 24h. Cells were washed and incubated at 37°C in the presence of 5% CO_2_ for 50 min with a hypotonic buffer. The supernatant from the cells was removed to a final 21 μl volume and 25 μl of compound solution diluted in rubidium containing buffer was added. After 10 min incubation, cells were washed with a hypotonic buffer and 21 μl of 2% tergitol NP-40 was added per well. Plates were sealed and stored overnight at room temperature. Finally, 30 μl of supernatant for each well were diluted in 170 μl of water and the rubidium content was measured using an atomic absorption spectrophotometer at 780 nm (Aurora Biomed, ICR 12000). Bumetanide was used as the reference compound for calculation of percentage of inhibition for each compound.

#### Morphology and cytotoxicity cellular assay

Cytotoxicity evaluation was performed on A549 human lung carcinoma cells (ATCC), employing a modified MTT (3-(4,5-dimethylthiazol-2-yl)-2,5-diphenyltetrazolium bromide) assay^46^. Indirect immunofluorescence was performed in A549 cells, as previously described^47^. An absorbance spectrum in methanol was made for each compound in a spectrophotometer Evolution 201 (Thermo Scientific) in order to identify the absorbance maxima.

#### Tubulin polymerization assay

The polymerization of 25 M tubulin in glycerol-assembling buffer, GAB (3.4 M glycerol, 10 mM sodium phosphate, 1 mM EGTA, 1 mM GTP, pH 6.5), in the presence of 27.5 M of the ligand studied (colchicine and compounds CT1, CT2, CT3, CT4 and CT5) or 2.75 l of DMSO (vehicle) was monitored along time by turbidity using a Varioskan Flash multimode microplate reader (Thermo Scientific) at a wavelength of 350 nm and 37 °C.

#### Tubulin binding assays

The effect of compounds CT1 and CT4 in the binding of 2-methoxy-5-(2,3,4-trimethoxyphenyl)-2,4,6-cycloheptatrien-1-one MTC^48^ was studied as previously described^49,50^. The effect of compound CT3 in the binding of (R)-(+)-ethyl 5-amino 2-methyl-1,2-dihydro-3-phenylpyrido[3,4-b]pyrazin-7-yl carbamate (R-PT), a well characterized reversible colchicine-binding were performed as described^51^. The binding of CT5 at the eribulin site was studied by monitoring the influence of eribulin in the kinetics of binding of the compound to tubulin. The compound undergoes a large increase in fluorescence at 465 nm (excitation 357 nm) upon incubation with tubulin. Their kinetics of binding of the compound to 10 M tubulin in NaPi buffer (10 mM Phosphate Buffer pH 7.0, 0.1 mM GTP) was measured in a Fluoromax-2 fluorimeter (Horiba) in the presence or absence of 50 M of vincristine, podophyllotoxin, maytansine or eribulin.

#### Molecular candidates for phenotypic screening

6 different tubulin binders (i.e. colchicine, paclitaxel, vinblastine, vincristine, vinpocetine and dolastatin 12) were used as reference compounds to search in the HTSFP PubChem collection with a Z_NMI_ threshold of 4.0. The resulting 1,559 molecules were structurally clustered using the ECFP4 fingerprints generated by Pipeline Pilot 9.2 and sorted according to Z_AG_. Known tubulin inhibitors and their chemically similar cluster members were removed and first 20 compounds were selected for confirmation.

#### Target identification assays

A detailed list of the assay conditions, detection methods and reference compounds for the different biochemical and cell-based assay employed in this work can be found in the Supplementary Table 4.

#### Code availability

Computer code for the BIOSEA implementation and HTSFP public data set are available for download from the authors’ repository at https://github.com/accsc/BIOSEA.

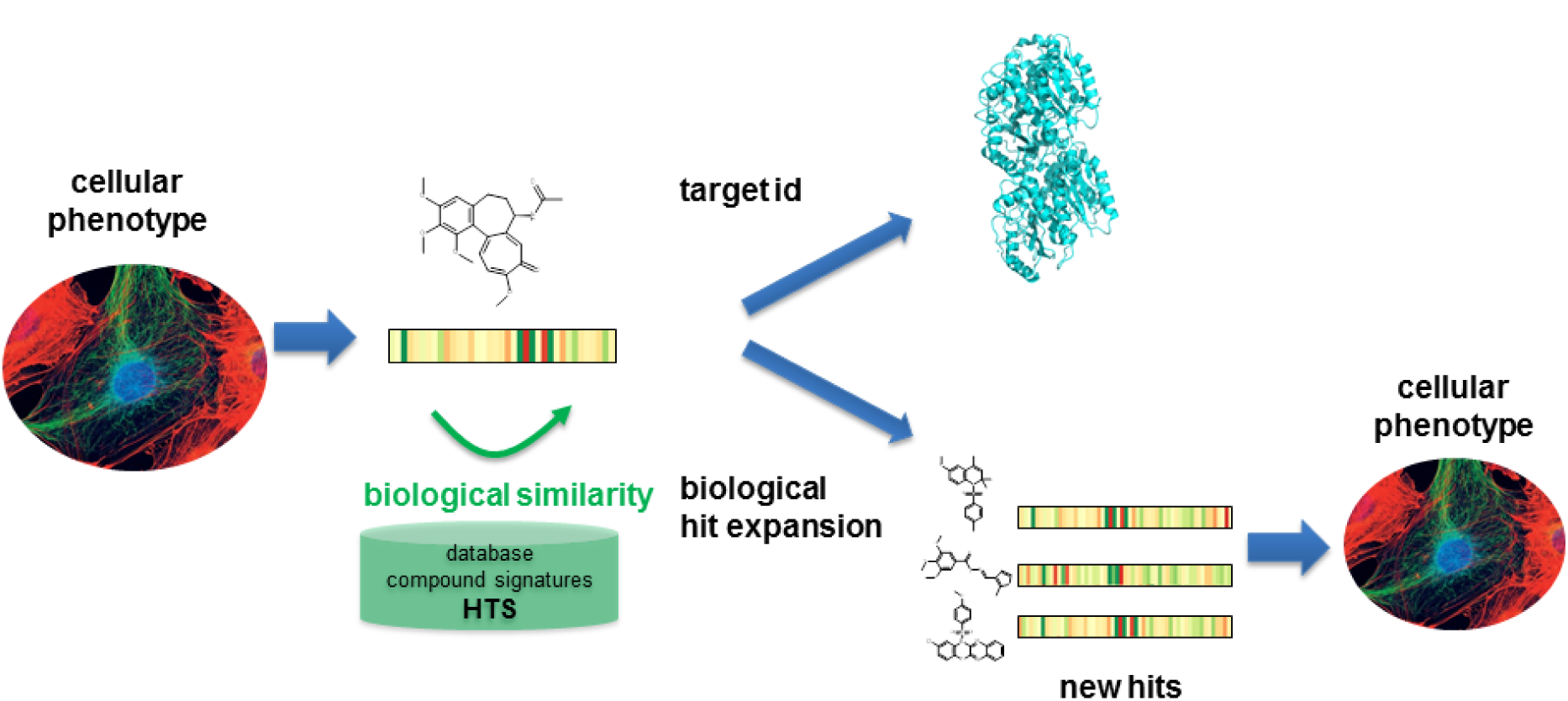
Graphical abstract.

## ACKNOWLEDGEMENTS

A.C. gratefully acknowledges the Roche Postdoctoral Fellowship Program for financial support. The authors thank specially Dr. Martin Erkens, Dr. Rene Wyler, Dr. Thilo Enderle, Dr. Lisa Sach-Peltason, Dr Samuel Croset and Dr. Oliv Eidam for their help and thoughtful comments and to Dr. Klaus Muller for his support and invaluable insight. Eurofins’ (CEREP) is gratefully acknowledged for providing us with single concentration activities from the BioPrint database in advantageous conditions.

## Author contributions

A.C.C. and P.P. designed the BIOSEA method and planned the target identification experiments. D.L, M. R., I.B. and F.D. designed and performed the cell cycle arrest phenotypic screening, the tubulin polymerization assay and the tubulin binding competition experiments. B.F. was the NKCC1 project leader. A.C., P.P, B.F. designed the NKCC1 hit expansion campaign.

